# Mechanistic picture for chemo-mechanical couplings in a bacterial proton-coupled oligopeptide transporter from Streptococcus thermophilus

**DOI:** 10.1101/2020.10.22.351015

**Authors:** Kalyan Immadisetty, Mahmoud Moradi

## Abstract

Proton-coupled oligopeptide transporters (POTs) use the proton electrochemical gradient to transport peptides across the cell membrane. Despite the significant biological and biomedical relevance of these proteins, a detailed mechanistic picture for chemo-mechanical couplings involved in substrate/proton transport and protein structural changes is missing. We therefore performed microsecond-level molecular dynamics (MD) simulations of bacterial POT transporter PepT_St_, which shares ~80% sequence identity with the human POT, PepT1, in the substrate binding region. Three different conformational states of PepT_St_ were simulated including, (i) occluded, *apo*, (ii) inward-facing, *apo*, and (iii) inward-facing_occluded_, Leu-Ala bound. We propose that the interaction of R33 with E299 and E300 acts as a conformational switch (i.e., to trigger the conformational change from an inward-to outward-facing state) in the substrate transport. Additionally, E299 and E400 should disengage from interacting with the substrate either through protonation or through co-ordination with a cation for the substrate to get transported.

## Introduction

Proton-coupled oligopeptide transporters (POTs) are membrane proteins present in the brush borders of kidneys and small intestine.^1^ They belong to the major facilitator super family (MFS) and important features of this family of proteins are substrate promiscuity and presence of multiple substrate binding regions,^2^ which are thought to be responsible for the ability of substrates/peptides to bind in multiple orientations ^1,3,4^ and with varying stoichiometry.^5^ This feature of POTs is of great interest due to its biomedical relevance.^6–8^ Human POTs (PepT1 and PepT2) absorb and retain a wide range of di- and tri-peptides and peptide like compounds. ^4,9^ They also recognize and help improve the bio-availability of poorly absorbed peptide-like drug compounds such as *β*-lactam antibiotics,^10^ angiotensin-converting enzyme inhibitors, and antiviral prodrugs. ^11–15^ Despite their biomedical relevance, several aspects of POTs are still debated, mainly substrate/peptide binding and transport mechanism and proton modulation of this process. This is mainly due to the non-availability of human POTs (PepT1 and PepT2) crystal structures. However, crystal structures of several bacterial homologues of human POTs, such as PepT_So_,^3,16^ PepT_St_,^17,18^ PepT_So2_,^7^ and GkPOT,^19^ are available.

It has been proposed that the POT transporters follow alternating access mechanism^20^ for transporting the substrates. ^17^ According to this mechanism, a POT transporter has to undergo at least four major conformational changes for it to facilitate transport, mainly outward-facing (apo), occluded (substrate bound), inward-facing (apo), and occluded (apo). ^1,4^ All the bacterial POT crystal structures reported to date are either occluded (OCC), ^18^ inward-facing_occluded_(IF_occ_) or an inward-facing (IF) state;^3,7,16–18,21,22^ an outward-facing (OF) state is not yet available. Even the MD studies that were reported earlier on POTs ^19,23,24^ failed to capture the fully open and stable OF conformation. Fowler et al. generated PepT_St_ model in an OF state through repeat-swapping technique, ^25,26^ however, they were not able to validate it through the DEER experiments.^24^

PepT_St_ is a bacterial POT symporter from Streptococcus thermophilus. Several studies have been carried to investigate different aspects of PepT_St_ through various experimental^5,18,22,24,27,28^ and computational techniques, ^24,28,29^ including solving protein structures in various conformations,^17,18,22,27^ understanding and proposing the functional mechanism,^4,17,24^ identifying different substrate binding sites and multiple binding modes for the substrates, ^4,5^ multispecific substrate recognition pathways,^3,18^ mechanism of proton and peptide coupling, ^28^ different transport mechanisms for different POT transporters, ^4,28^ and binding of a range of substrates. ^29^ However, a study utilizing all available conformations of the PepT_St_ is needed to gain a detailed atomistic level understanding of its transport mechanism and the current availability of multiple conformations of PepT_St_ provides a perfect opportunity.^17,18,22,27^

PepT_St_ is the only POT symporter that has been crystallized in OCC, IF, and IF_occ_ conformational states. ^17,18,22,27^ Hence, in this study we simulated all three available conformational states of PepT_St_ via unbiased all-atom MD^30–33^ with either E300 protonated or deprotonated. Additionally, we simulated the IF_occ_(LA) in all possible protonation combinations of E299, E300, and E400 residues, which resulted in a total of eight substrate bound systems (see methods for more details). Substrate LA binds with millimolar affinity (≈0.56 mM) to PepT_St_.^18^ It was demonstrated that the three glutamates lining the substrate translocation path (i.e., E299, E300 and E400) are very key for functioning of the PepT_St_,^4,17,18^ and E400 in particular is conserved across all members of the family.

Overall, we simulated 12 different systems (see Table 1 for details), each for ≈1*μs* (12 *μ*s in total and this is the highest simulation time for any POT transporter reported thus far). We propose that (1) the R33-E299 and the R33-E300 salt bridge interactions act as a molecular switch for the protein to undergo complete conformational change from an IF to OF side, (2) deprotonation of the E300 and thus the formation of R33-E300 salt bridge is crucial for the protein to transform from OF to occluded/IF side, (3) disengaging of E299 and E400 from interacting with the substrate LA is key for it to get transported, (4) either protonation of E299 and/or E400 or co-ordination of E299 and E400 with sodium prevent these residues to engage with the substrate and facilitate its transport, (5) the C-domain is more dynamic than the N-domain and the C-domain plays a key role in the transportation process, particularly TMs 7, 10, and 11.

**Table 1:** Different systems simulated via MD.

| System | UP:E300 | P:E300 | P:E299 | P:E400 | P:E299,E300 | P:E299,E400 | P:E300,E400 | P:E299,E300,E400 |
| --- | --- | --- | --- | --- | --- | --- | --- | --- |
| Occ(Apo)<sup>a</sup> | 1μs | 1μs | | | | | | |
| IF(Apo)<sup>b</sup> | 1μs | 1μs | | | | | | |
| IF<sub>Occ</sub>(LA)<sup>c</sup> | 1μs | 1μs | 1μs | 1μs | 1μs | 1μs | 1μs | 1μs |

<sup>a</sup> Occluded PepT<sub>St</sub> in apo form; <sup>b</sup> Inward-facing PepT<sub>St</sub> in apo form. <sup>c</sup> Inward-facing Occluded PepT<sub>St</sub> bound with substrate LA. UP and P refers to deprotonated and protonated residues.

## Results and discussion

### R33-E300 and R33-E299 salt bridge interactions act as a conformational switch for PepTSt to transition from an IF(Apo) to OF(Apo) state

E300 belongs to TM7 and reside at the intersection of C- and N-domains. Studies reported that E300 plays a key role in functional dynamics of the POT family of transporters by forming a salt bridge with R33 (that belongs to TM1) and stabilizing the IF conformation, at least in GkPOT and PepT_St_.^1,17,19^ It was also identified that R33 or its equivalent is key for proton-coupled transport in the GkPOT and PepT_St_.^19^ The R33-E300 salt bridge forms upon deprotonation of the E300 bringing the two domains closer, and breaks when the E300 is protonated separating the two domains. ^1,19^ Therefore, protonation of the E300 and subsequent breaking of the R33-E300 salt bridge is required for the protein transitioning from an IF/IF_occ_ to OF state, which facilitates substrate entering the protein. To test this hypothesis we measured the R33-E300 salt bridge distance.

Our simulation data suggests that the R33-E300 salt bridge forms in PepT_St_ upon de-protonation of the E300 as expected (Fig. 1E and Fig. S1). In both the E300 deprotonated apo systems (i.e., UP:E300:OCC(Apo) and UP:E300:IF(Apo)) the R33-E300 salt bridge stabilized around 3 Å (Fig. 1E and Fig. S1 B,C). However, this salt bridge broke in the two apo E300 protonated systems (i.e., P:E300:OCC(Apo) and P:E300:IF(Apo)), and the distance fluctuated around 5 - 7Å (Fig. 1E and Fig. S1 E,F). In the other four E300 protonated systems with substrate LA bound (i.e., P:E300:IF_occ_(LA), P:E299,E300:IF_occ_(LA), P:E300,E400:IF_occ_(LA), and P:E299,E300,E400:IF_occ_(LA)), the R33-E300 salt bridge was broken as expected and the distances varied between 7 - 11Å (Fig. 1E and Fig. S1 D, I, K and L). Particularly, in the P:E300:IF_occ_(LA) system, it went up to 11Å (Fig. 1E and Fig. S1 D); substrate LA was situated between the two domains and interacting with R33 and E299 (as shown in the contact analysis (Table. S1)). *This (P:E300:IF*_*occ*_*(LA)) probably represents the OF state of the PepT*_*St*_ (Fig. S6A, right panel). Also, this probably is the first step for the entry of the substrate into the substrate translocation site. In the two E300 protonated apo systems (i.e., P:E300:OCC(Apo) and P:E300:IF(Apo)), although the R33-E300 was broken, R33 was engaging in a salt bridge formation with the E299 preventing transitioning of the protein completely from an IF/IF_occ_ to OF state (Fig. 1F and Fig. S2 E, F). Of note, the R33-E299 salt bridge was absent in the remaining 10 systems. *Hence, we propose that for this protein to completely transition from the IF(apo) to the OF(apo) side, both E299 and E300 have to be protonated. Also, we propose that the two salt bridges (i.e., R33-E300 and R33-E299) act as a conformational switch for PepT*_*St*_ *to transition from the IF(Apo) to the OF(Apo) state.* In the four other systems with E300 deprotonated and LA bound (i.e., UP:E300:IF_occ_(LA), P:E299:IF_occ_(LA), P:E400:IF_occ_(LA) and P:E299,E400:IF_occ_(LA)), R33-E300 salt bridge was intact, although fluctuating often (Fig. 1E and Fig. S1 A, G, J and H). This situation forced the substrate LA towards the cytoplasmic side, therefore, in the process of attaining a new optimal orientation/conformation substrate LA was destabilizing the protein, particularly the R33-E300 salt bridge, because it often interacts with the E300 and destabilizing the R33-E300 salt bridge (Table. S1). When E299 or E400 were deprotonated, substrate LA moved down the binding site and interacted with these two residues; therefore, it is less likely that the substrate LA disturbs the R33-E300 salt bridge in these two systems. This was reflected in the P:E299:IF_occ_(LA) and the P:E400:IF_occ_(LA) systems; in the first case LA was interacting with the E400 (≈ 98% contact frequency) and in the second case it was interacting with the E299 (≈ 90% contact frequency) (Table. S1). In either case, E300 was not among the top ten interacting residues with LA (Table. S1). When E299 and/or E400 were not available, either due to protonation or for other reasons, LA was engaging with the E300. This was reflected in the P:E299,E400:IF_occ_(LA) and UP:E300:IF_occ_(LA) systems. In either case, LA was interacting with the E300 for at least 30% of the simulated time (Table. S1). Overall, the four E300 deprotonated holo systems demonstrated that the substrate LA was not comfortable in the binding site when protein was in the IF_occ_ state, hence, it facilitates transitioning of the protein completely towards the cytoplasmic side, which in-turn facilitates the substrate transport.

**Figure 1:**
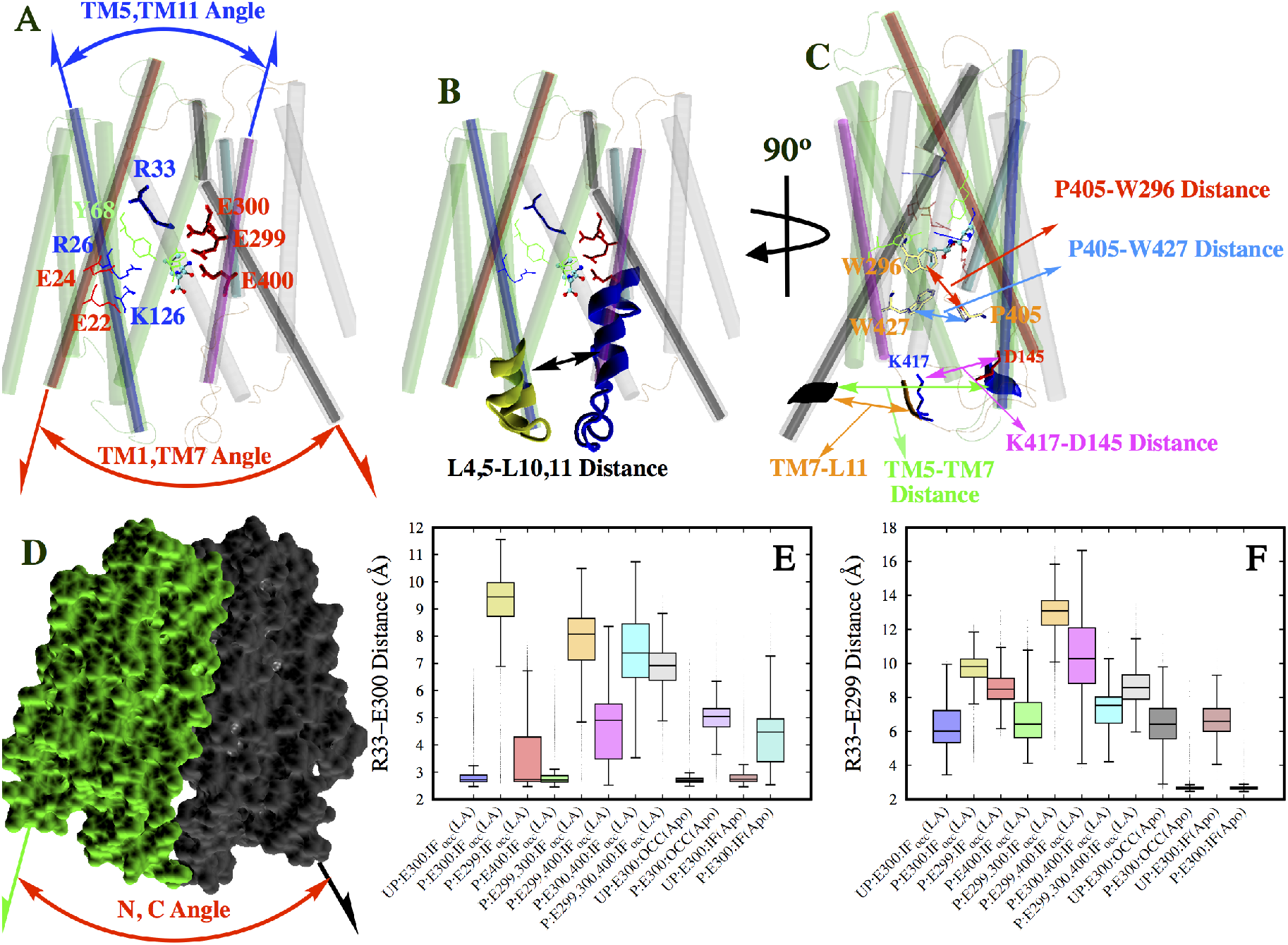
**A** PepT_St_ comprises 14 transmembrane helices (TMs), 12 are arranged in two bundles as N- (colored green, TMs 1-6) and C-domains (colored grey, TMs 7-12). The two other helices are not shown since their role in functioning of the POTs is not yet established. Positive and negative residues are colored blue and red, respectively. Substrate LA that is binding between the two domains is colored cyan and shown in ball and sticks representation. Inter-helical angles TM1-TM7 (colored red) and TM5-TM11 (colored blue) represent extracellular and intracellular gates, respectively, which are thought to control the substrate transport.^1,4^ **B** Distance between L4,5 and L10,11. L4,5 is the loop between the helices TM4 and TM5. L10,11 is the loop between the helices TM10 and TM11. **C** Various other variables measured along the substrate translocation path. **D** N-,C-interdomain angle. N- and C-terminal domains are colored green and orange, respectively. **E, F** Box plots of distances of R33-E300 and R33-E299 salt bridges. The presence of substrate LA destabilized R33-E300 salt bridge in all systems with the E300 deprotonated, where as the absence of it keeps this salt bridge intact. This salt bridge is lost in all the E300 deprotonated systems as expected, although in the apo systems this salt bridge distance is smaller compared to the substrate bound. The R33-E299 salt bridge, except in the P:E300:OCC(Apo) and P:E300:IF(Apo) systems, is absent in all other systems.

### Deprotonation of E300 is required for the substrate to get transported towards the cytoplasmic side

Of the eight holo systems, in half of them E300 was deprotonated(i.e., UP:E300:IF_occ_(LA), P:E299:IF_occ_(LA), P:E400:IF_occ_(LA), P:E299,E400:IF_occ_(LA)), and in these four systems substrate moved towards the cytoplasmic side (Fig. 2B), since the N- and C-domains in the OF side are locked by the R33-E300 salt bridge. However, in the four E300 protonated holo systems (i.e., P:E300:IF_occ_(LA), P:E299,E300:IF_occ_(LA), P:E300,E400:IF_occ_(LA) and P:E299,E300,E400:IF_occ_(LA)), substrate stayed more towards the OF side compared to the E300 deprotonated systems; the difference in the substrate Z-values (a variable we defined to describe the position of the substrate along the membrane normal) between the E300 deprotonated/protonated systems is statistically significant and they do not overlap (Fig. 2B). This is because the R33-E300 salt bridge was broken in these systems (Fig. 1E) providing more room for the substrate to move up the binding site and attain an ideal orientation. This was also reflected in the substrate overall RMSD (Fig. 2A). Although, there was a sharp increase in the four holo E300 protonated systems in the beginning (Fig. S3 D, I, K and L), later they were overall stable compared to the four holo E300 deprotonated systems (Fig. S3 A, G, H and J). Substrate overall RMSD is the relative change in the substrate RMSD with respect to its starting conformation, and this was calculated by aligning the protein against its own starting conformation. ^1^ On the other hand, internal RMSD depicts the conformational changes happening in the substrate, and was measured by aligning the substrate against its own initial orientation. Internal RMSDs of substrates were fairly stable in all eight holo systems studied (fluctuated around 2 Å(Fig. 2C)), hinting that there were no significant conformational changes in the substrate.

**Figure 2:**
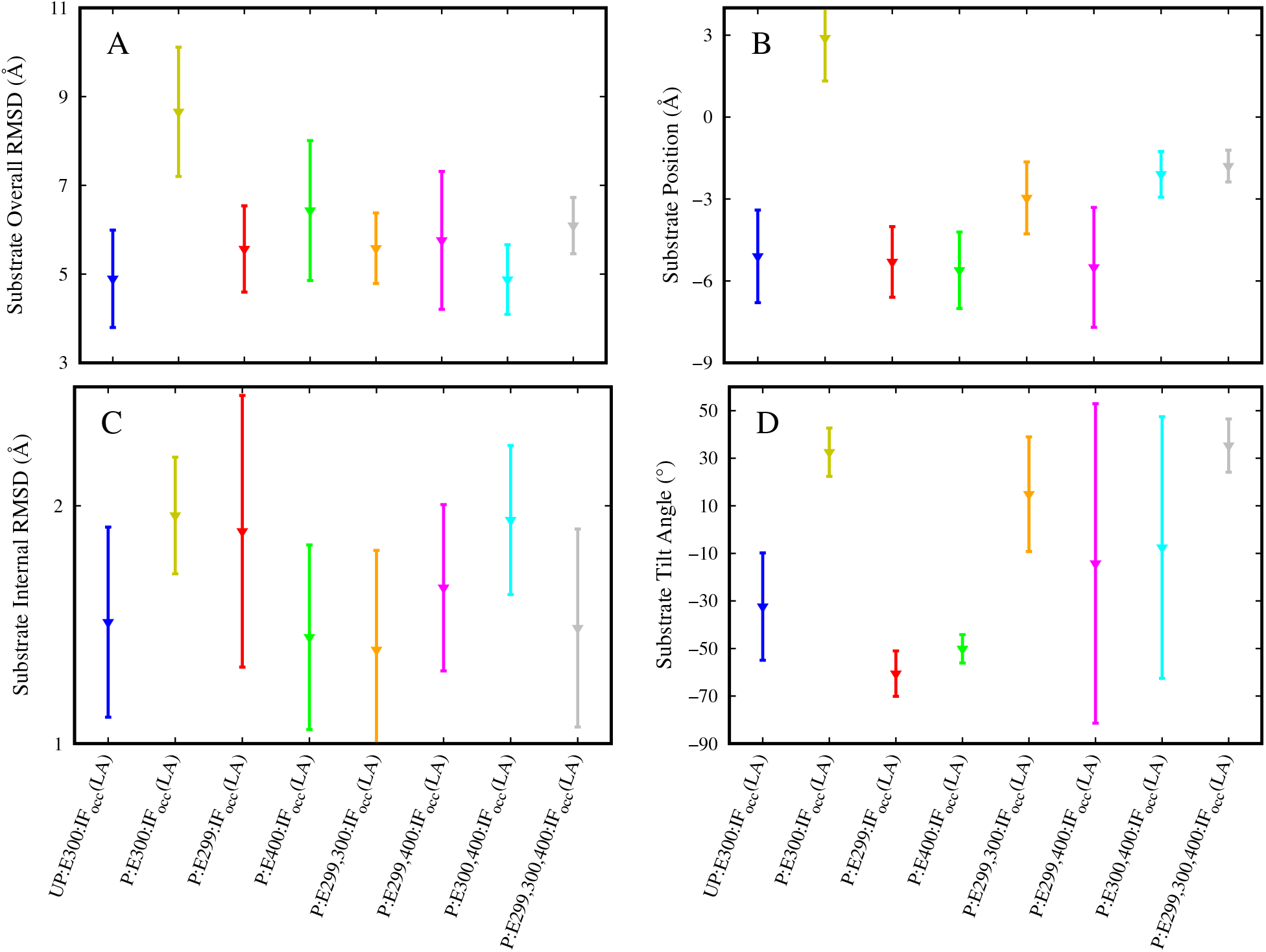
Effect of deprotonation/protonation of residues on the substrate overall (**A**) and internal (**C**) RMSDs, substrate position (**B**) along membrane normal and substrate tilt angle (**D**). In the P:E300:IF_occ_(LA) system, substrate moved up in the binding site, which is not seen in other systems (**B**). The overall RMSD of substrate in the P:E300:IF_occ_(LA) system was relatively higher compared to other seven systems, which indicates that E300 is the most sensitive and important residue that plays a key role in PepT_St_ dynamics (**A**).

Among the four E300 deprotonated holo systems, substrate was completely transported in two systems (i.e., inUP:E300:IF_occ_(LA) and P:E400:IF_occ_(LA)) towards the IF side, as reflected in the sudden drop in substrate Z values Fig. 3F; and in the other two holo systems (i.e., P:E299:IF_occ_(LA) and P:E299,E400:IF_occ_(LA)) substrate translocated towards the IF side waiting to get transported (Fig. 2B). In the UP:E300:IF_occ_(LA) and P:E400:IF_occ_(LA) systems, substrate attained a vertical orientation (Fig. 3 I,L) and then left the protein towards the IF side around 900 and 990 ns, respectively (Fig. 3F), confirming that these were not favorable conditions for the binding of the substrate LA in the binding site and thus eventually got transported.

**Figure 3:**
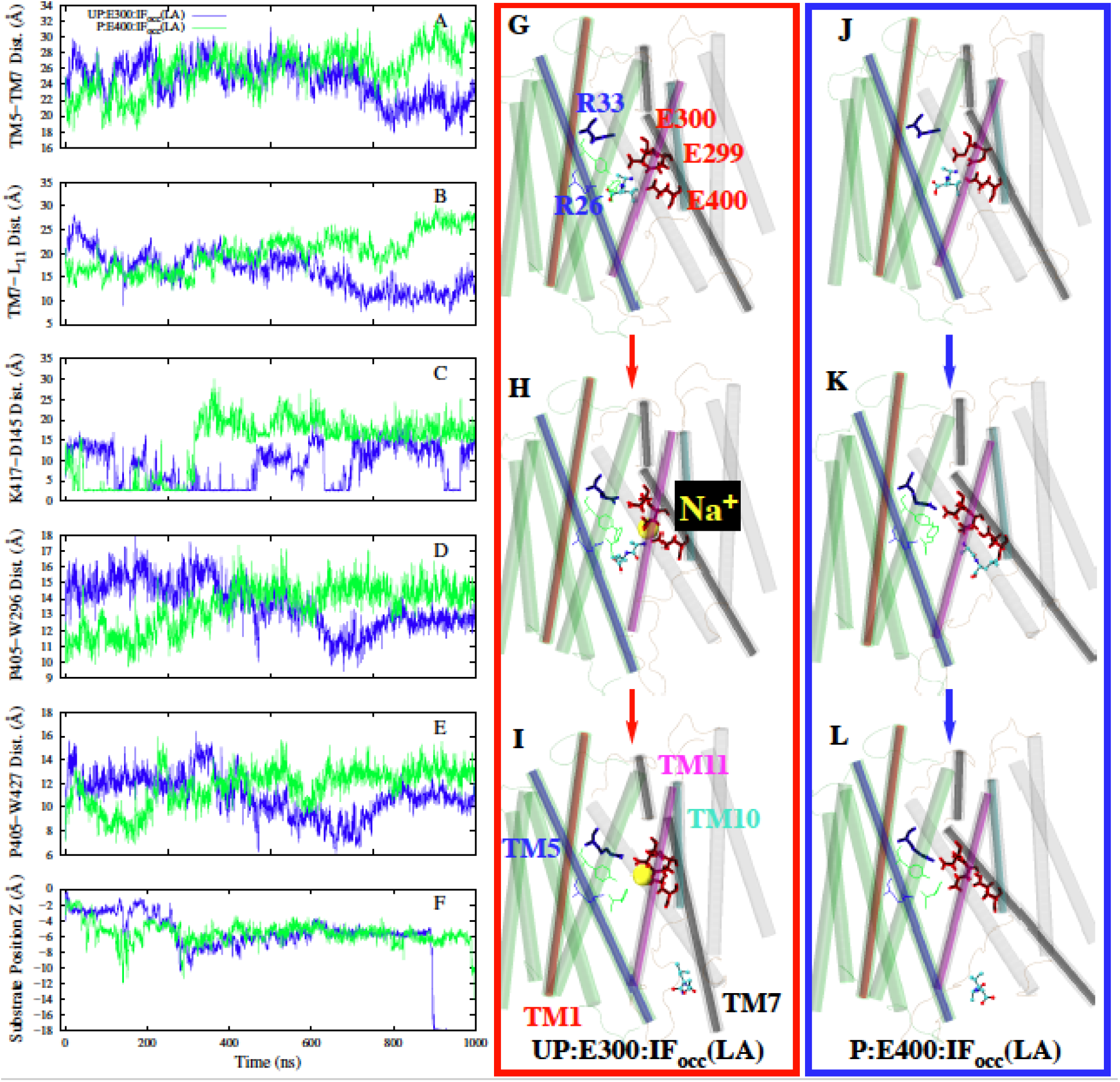
Pathway of substrate release. Shown in left panels are various variables measured across the substrate transport path (**A-F**). Middle panels (**G-I**) and right panels (**J-L**) shows the transportation of substrate LA in the UP:E300:IF_occ_(LA) and in the P:E400:IF_occ_(LA) systems, respectively. All the variables were explained in Fig. 1.

To assess orientation of the substrate LA, we measured tilt angle, which describes deviation of the substrate from membrane normal. Tilt angle zero indicates that the substrate attained a horizontal orientation, where as negative or positive tilt angles infer that the substrate attained a vertical orientation. Also, negative tilt angle indicates that the C- and N-terminus faces cytoplasm and periplasm, respectively, and positive tilt angle indicates the opposite. We have observed a correlation between the substrate tilt angle and protonation of E300. When E300 was protonated, tilt angle was positive (i.e., C-terminus was facing the periplasmic side), and vice versa (i.e., C-terminus was facing cytoplasmic side) (Fig. 2D); the P:E299,E400:IF_occ_(LA) and P:E300,E400:IF_occ_(LA) systems were exceptions, where they undergo a range of fluctuations, as reflected by the huge standard deviations. Overall, we show that for the substrate to bind comfortably in the pocket E300 has to be protonated. However, for the substrate to get transported, formation of the R33-E300 salt bridge is mandatory, which in turn requires deprotonation of E300. When the R33-E300 salt bridge was formed, it pushes the substrate towards the cytoplasmic side leaving less room for the substrate to adjust to the binding site. Also, the R33-E300 salt bridge formation opens the protein towards the IF side (discussed in detail in the next section), exposing the substrate to water and eventually getting transported towards the cytoplasmic side.

### Disengaging of E299 and E400 with the substrate is key for the substrate to release into the IF side

We have observed that E299 and E400 were playing a key role in the transport of the substrate LA. These two residues act as the primary interaction sites for the substrate (Table. S1) after it has been pushed down following deprotonation of E300 and the subsequent formation of the R33-E300 salt bridge, hence preventing the transport of the substrate towards the cytoplasmic side, at least temporarily. We identified two events that could circumvent this problem by disrupting interaction of the residues E299 and E400 with the substrate LA and thus promote its transport: (1) co-ordination of the residues E299 and E400 by sodium ions and (2) protonation of E299 or E400 or both. Both events are discussed in detail below.

#### Sodium modulation of substrate binding and transport

Several membrane transporters rely on ions for transporting the substrate across the membrane bilayer, for example the sodium-coupled transporters.^30^ In this study we observed that sodium ions play a role in the substrate LA transport. Of the 12 simulated systems, in five of them we have observed at least one sodium ion co-coordinating the E299 and E400 residues; two were holo (UP:E300:IF_occ_(LA) and P:E300:IF_occ_(LA)) and the remaining three were apo (UP:E300:OCC(Apo), UP:E300:IF(Apo) and P:E300:IF(Apo)) (Fig. 4A). On a side note, for the sodium co-ordination to happen both the residues E299 and E400 have to be deprotonated, which is the case with the five systems mentioned previously.

**Figure 4:**
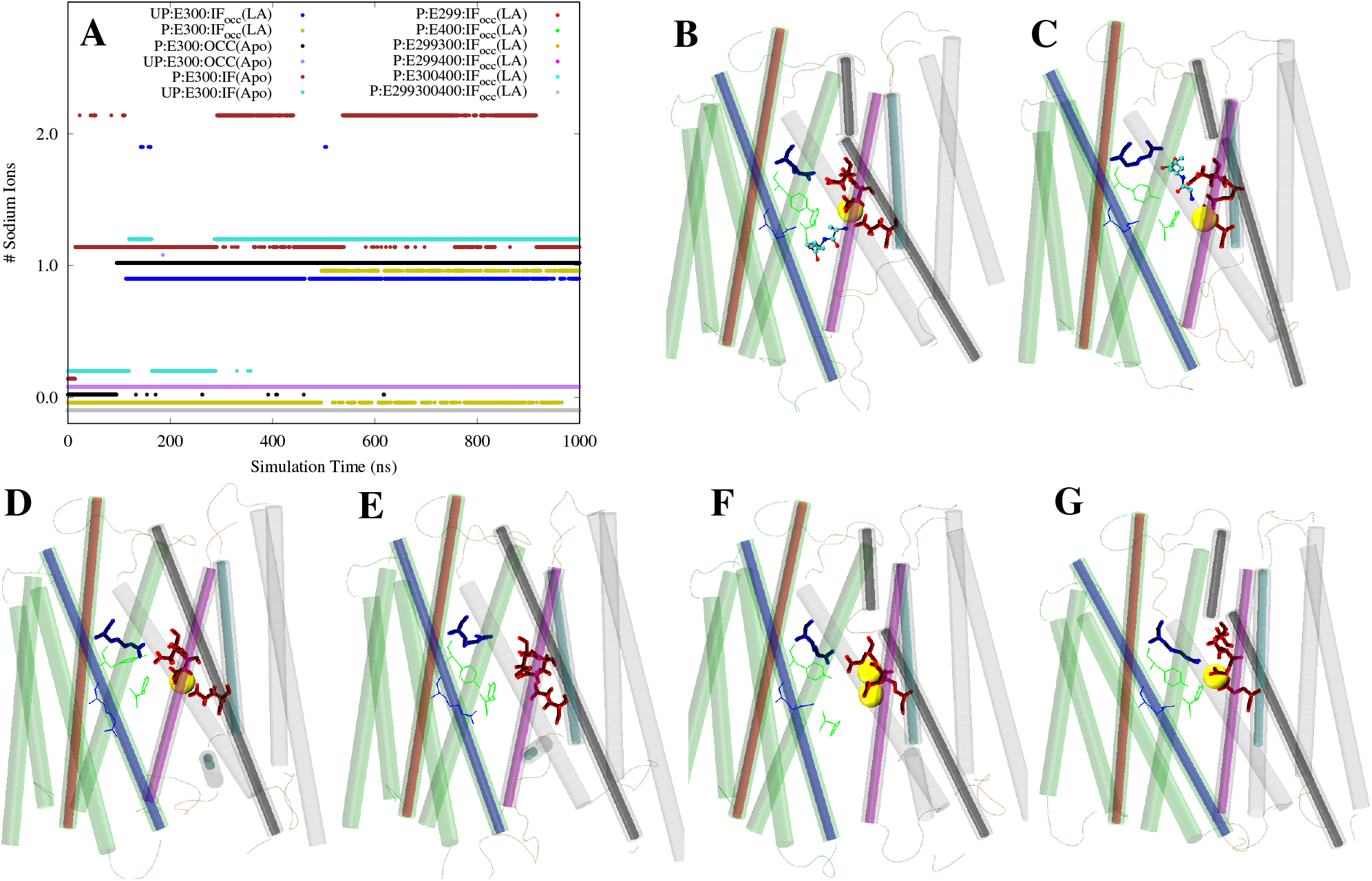
Sodium co-ordination of E299 and E400. Sodium ions within 4 Å of both 299 and E400 are shown. **A** is time series of sodium co-ordination. **B** is UP:E300:IF_occ_(LA), **C** is P:E300:IF_occ_(LA), **D** is UP:E300:OCC(Apo), **E** is P:E300:OCC(Apo), **F** is UP:E300:IF(Apo), and **G** is P:E300:IF(Apo). Sodium ions are represented as yellow spheres. Sodium co-ordination was not seen in the case of P:E300:IF_occ_(LA).

In the UP:E300:IF_occ_(LA) system, E299 and E400 were coordinated by sodium preventing them from engaging with the substrate LA (Table. S1), thus facilitating the substrate transport. Additionally, in the UP:E300:IF_occ_(LA) system E300 was deprotonated, therefore the R33-E300 salt bridge was intact, which pushed the substrate down and contributed to substrate transport. In the other substrate bound system (i.e., the P:E300:IF_occ_(LA)), although E299 and E400 were deprotonated and co-ordinated by the sodium, substrate was not transported. This may be because in this system E300 was protonated, therefore the R33-E300 salt bridge was not intact providing more room for the substrate to bind com-fortably in the binding site between the residues R33 and E299. *Hence, we propose that deprotonation of E300 is not just enough for the transportation of the substrate, but also the residues E299 and/or E400 should be disengaged with the substrate. One way of achieving this is through sodium co-ordination of the residues E299 and E400, which is only possible when E299 and E400 were deprotonated as discussed earlier.* In the other seven systems in which sodium co-ordination of E299 and E400 was not observed, either E299 or E400 or both were protonated hindering the co-ordination.

#### Protonation of E299/E400 promoting transport

We have demonstrated thus far that for the substrate to get transported towards the IF side the R33-E300 salt bridge has to be intact, which is only possible when the E300 is deprotonated. Our simulated set contains four E300 deprotonated holo systems and in three of them either E299 and/or E400 were protonated(i.e., P:E299:IF_occ_(LA), P:E400:IF_occ_(LA) and P:E299,E400:IF_occ_(LA)). We previously showed that in the UP:E300:IF_occ_(LA) system substrate got transported towards the IF side. Also, in the P:E400:IF_occ_(LA) system substrate completely left the pocket (Fig. 3F). In the remaining two systems (i.e., in P:E299:IF_occ_(LA) and P:E299,E400:IF_occ_(LA)), substrate moved down the binding site and was about to leave as reflected by the substrate Z values (Fig. 2B). In the P:E299:IF_occ_(LA) system, although E299 was protonated, substrate did not leave in a 1*μs* simulation since E400 was interacting with substrate and preventing it from leaving as shown in contact frequency analysis (Table S1). Lastly, in the P:E299,E400:IF_occ_(LA) system, substrate was engaging with the residues R26, Y68, and E300 (Table S1), hence was not transported. This was also reflected in the R30-E300 salt bridge distance; substrate LA disrupted this salt bridge (Fig. S1 J) while interacting with the residue E300 (Table S1). We demonstrated here that deprotonation of E300 and protonation of E299/E400 facilitates transport, since the protonated E299/E400 disengages themselves from the substrate.

### Ideal conditions for the substrate to get transported

Based on the eight holo systems studied here, systems that are ideal for the substrate to get transported are: (1) UP:E300:IF_occ_(LA), (2) P:E400:IF_occ_(LA), (3) P:E299:IF_occ_(LA) and (4) P:E299,E400:IF_occ_(LA). The common factor among these four systems is that E300 is deprotonated, hence, the R33-E300 salt bridge was intact (Fig: 1 E,F and Fig. S1) in all four systems. Therefore, substrate was pushed down the translocation path waiting to get transported (Fig. 2B), which establishes that the R33-E300 salt bridge formation is key for it to get released. In the first case (i.e., the UP:E300:IF_occ_(LA)), co-ordination of E299/E400 with a sodium ion disengages substrate from interacting with the residues E299 and E400 (Fig. 4A), facilitating the substrate release. In case 2 (i.e., the P:E400:IF_occ_(LA)), since E400 was protonated it was not engaged with the substrate. Although, E299 was deprotonated and engaging with the substrate blocking the release (Table. S1, 90% contact frequency), since E299 was stacked between the residues E300 and E400 it cannot hold the substrate for long, facilitating the substrate release. In case 3 (i.e., the P:E299:IF_occ_(LA)), substrate moved down the pocket and is waiting to get released (Fig. 2B). In this system E400 (which is deprotonated) was engaging with the substrate (Table. S1, 98% contact frequency) impeding the release, where as the E299 cannot engage with the substrate since it was protonated. In case 4 (i.e., the P:E299:IF_occ_(LA)), both E299 and E400 were protonated, hence, cannot engage with the substrate. Instead, the substrate was engaging with the residues R26, Y68, and E300 with 50%, 66%, and 32% contact frequencies, respectively (Table. S1). The substrate moved down the binding site (Fig. 2B) and was visibly not stable as reflected in the overall RMSD (Fig. S3J) and waiting to get transported. *Overall, we show that the ideal scenario for the substrate to get transported is through the deprotonation of E300 and disengaging of E299 and/or E400 with the substrate (either through sodium co-ordination or protonation)*.

### Effect of deprotonation/protonation on protein global dynamics

We monitored the impact of deprotonation/protonation at the global level (i.e., on the entire protein). To asses the stability and also to identify major conformational changes we measured backbone RMSDs of protein. In all 12 cases protein reached equilibrium and RMSDs of all systems stabilized around 3 Å(Fig. S3). Protein RMSDs did not provide any clues of huge conformational changes at the global level.

Further, to asses conformational landscape of the 12 systems we employed principle component analysis (PCA).^31,34^ Impact of protonation/deprotonation on the protein conformation was clearly observed in the PCA analysis. When all 12 structures were projected on to the PC space, PC1 could clearly differentiate the entire trajectory into 2 distinguished clusters (Fig. 5A). Cluster one comprised of the two occluded(Apo) structures, and the second cluster consists of rest of the ten systems (i.e., the IF(Apo) and the IF_occ_ (LA bound)). Trajectories were clustered into three groups in the PC2 space, one cluster formed by the P:E300:IF(Apo), and second by the UP:E300:IF(Apo) and third by the rest. Square displacements have been calculated to identify the major contributions to the individual PC components. PC1 for the entire 12 system trajectory was dominated by the residues 163-164 (belongs to TM5) and 261 (belongs to a helix that is not part of the N- and C-domains) (Fig. 5D), where as PC2 was dominated by the residues 400-410 (loop between helices TM10 and TM11). PC1 and PC2 contributes ≈52% and ≈10% (Fig. 5G) to the total variance, respectively. When just substrate bound systems were projected on to the PC space, four clusters were observed; P:E400:IF_occ_(LA) was in the first cluster, UP:E300:IF_occ_(LA) in the second, P:E299,E300,E400:IF_occ_(LA) in the third, and the fourth cluster comprised of the rest (Fig. 5B). PC1 clearly differentiates P:E400:IF_occ_(LA) and UP:E300:IF_occ_(LA) structures when the substrates were released, indicating that these systems adopt different conformations when they release the substrates. Square displacements for just the substrate bound trajectories were dominated by C-domain in both the PC1 and PC2 modes (Fig. 5E). However, the first two PCs contributes only ≈30% to the total variation, which indicates that variation for this PC space was multifold and the first two PCs could not explain the entire story (Fig. 5H). When OCC(Apo), IF(Apo), UP:E300:IF_occ_(LA), P:E300:IF_occ_(LA), and P:E400:IF_occ_(LA) were projected together, both UP:E300:IF_occ_(LA) and P:E400:IF_occ_(LA) were clustered differently from the IF(Apo) systems on the PC2 space. This indicates that the UP:E300:IF_occ_(LA) and P:E400:IF_occ_(LA) structures were not as open as IF(Apo) systems when the substrate was released (Fig. 5C). Overall, PCA analysis highlights the sensitivity of POTs structure and conformational dynamics upon protonation/deprotonation of select few important residues. More importantly, this analysis also highlights that POTs can adopt different conformations while they release the substrate depending on the situation, which is also a main USP of the POT family of transporters. ^18^

**Figure 5:**
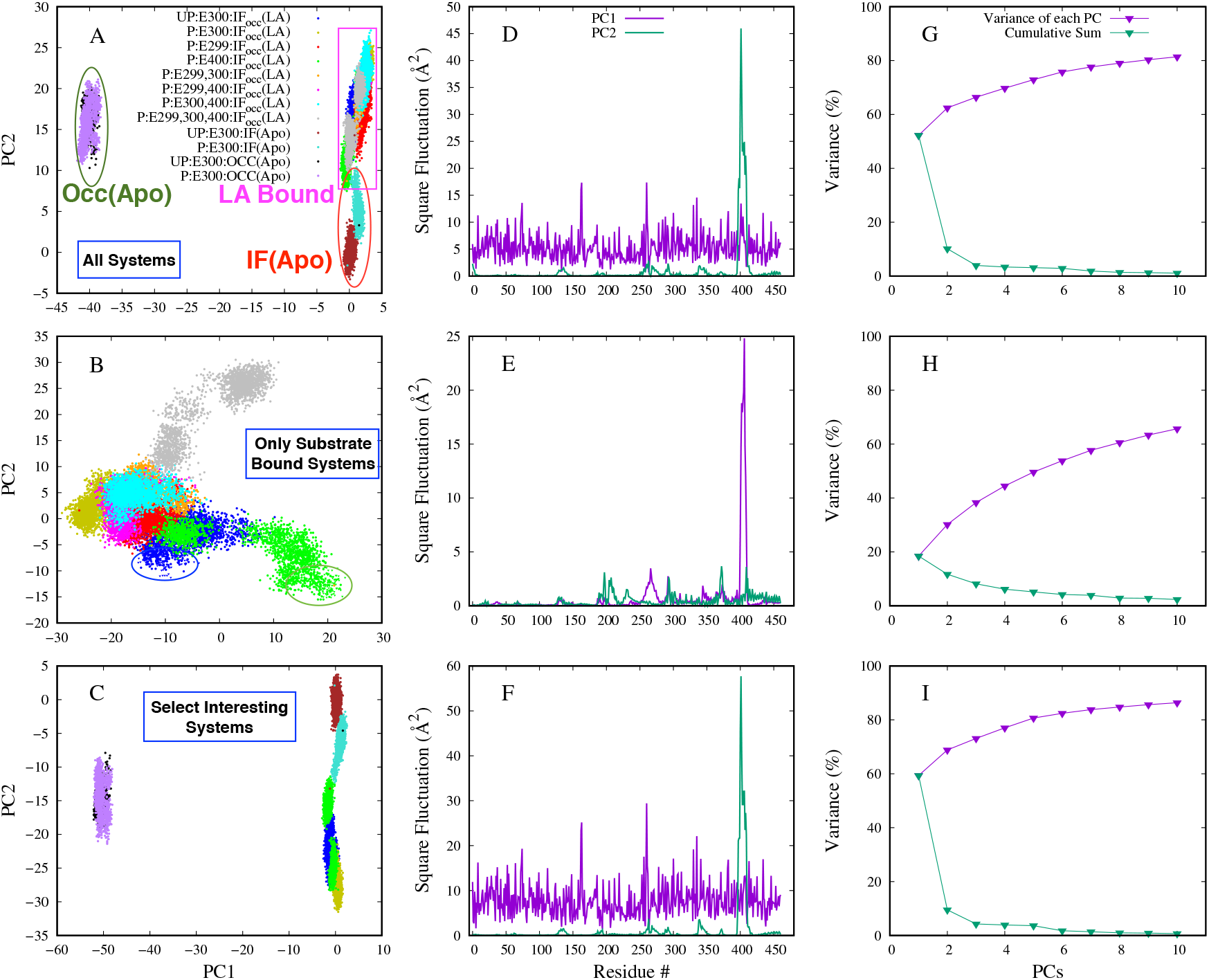
Principle component analysis. Left panels are the projection of data on to the PC1 and PC2 space. In **A** all systems were considered, in **B** just the substrate bound systems, and in **C** only select interesting systems were considered. Data points highlighted in circles (**B**) represent the protein conformations when the substrate left the protein into the cytoplasmic side. Middle panels (**D, E, F**) are square displacement of PC1 (purple) and PC2 (green) modes as a function of residue number. Right panels (**G, H, I**) depict the proportion of variance of the top ten modes in each case (green). Purple colored data points represent the cumulative average of the top ten PC modes.

Additionally, we calculated various variables to quantify global dynamics of the protein such as N-,C- inter-domain angle (Fig. 1D), TM1-TM7 and TM5-TM11 interhelical angles (Fig. 1A), and L_4,5_-L_10,11_ distance (Fig. 1B).^1^ L_4,5_ is a loop between the helices 4 and 5 and L_10,11_ is a loop between the helices 10 and 11 on the cytoplasmic side. L_4,5_-L_10,11_ distance quantifies opening of the protein on the cytoplasmic side. ^1^ According to this measure, the major impact of protonation was observed in the case of IF(Apo) systems (Fig. S6D); opening of the protein on the cytoplasmic side was greater in the UP:E300:IF(Apo) system compared to the P:E300:IF(Apo) (≈7Å) and the difference was statistically significant. Similar behavior was observed in the Occ(Apo) systems, although the difference between deprotonated and protonated systems was significant, the difference was not huge. Overall, in the apo systems (IF or OCC), E300 deprotonation makes the protein open more towards the cytoplasm than the E300 protonation systems. However, this trend was reversed in the holo systems, except in the case of P:E299:IF_occ_(LA). This behavior hints that the substrate binding favors more of an occluded conformation when E300 is protonated. The difference between different substrate bound systems is significant, although not huge. The N-,C- inter-domain angle indicates the opening between the two domains on the cytoplasmic side. The behavior of this variable was similar to the L_4,5_-L_10,11_ distance in the case of IF(apo) systems. The difference in N-,C- inter-domain angle between the UP:E300:IF(Apo) and P:E300:IF(Apo) systems was ≈10°; in the rest of the ten systems, no conclusive trend was observed. Similar behavior was observed in the case of TM1-TM7 (Fig. S6B) and TM5-TM11 (Fig. S6C) interhelical angles as well. TM1-TM7 interhelical angle is a hypothesized cytoplasmic gate, where as the TM5-TM11 interhelical angle is a hypothesized periplasmic gate.

### Pathway and mode of substrate transport

Based on the two substrate release simulations (i.e., UP:E300:IF_occ_(LA) and P:E400:IF_occ_(LA)) we mapped the substrate transport pathway in PepT_St_. We propose that this transition occurs in three stages.

In stage one, substrate undergoes orientation change, i.e., from horizontal to vertical (Fig. 3 H,K). In the vertical orientation the C-terminus and the N-terminus were facing the cytoplasmic and the periplasmic sides, respectively (which was also shown through the tilt angle (Fig. 2D)), since the substrate prefers to interact with the negatively charged residues (i.e., E299, E300, and E400) on the periplasmic side via the N-terminus. This behavior is very similar to the GkPOT.^1^ Note that the substrate is in a horizontal orientation in the crystal structures (Fig. 3 G,J).

In stage two, opening of the *hydrophobic bridge* facilitates downward movement of the substrate along the translocation path. This hydrophobic bridge is formed by the residues W296 (TM7), P405 (TM10) and W427 (TM11). These three residues belong to the C-domain, located in the middle of the protein right below the substrate binding site (Fig. 1C) and prevent the release of the substrate towards the IF side. Earlier it was reported that mutation of W427 to alanine drastically reduced the proton/peptide drive counter flow uptake.^17^ To map the opening and closing of this hydrophobic bridge we measured two different distances between: (1) the residues P405 and W427 and (2) the residues P405 and W296 (Fig. 1C). Both these distances increased in the course of simulation (Fig. 3D,E), which establishes that the opening of this hydrophobic gate is crucial for the transport of the substrate. However, the extent of the opening is greater in the case of P:E400:IF_occ_(LA) than in the P:E400:IF_occ_(LA).

In stage three, helices 5, 7, and 11 moved away from each other at the cytoplasmic side facilitating release of the substrate. To quantify this behavior we measured the distance between the tips of TM5 (considered residues 144, 145) and TM7 (considered residues 280, 281). Additionally, we measured distance between the tips of TM7 (considered residues 280,281) and TM11 (considered residues 418,419 that are part of the loop) (Fig. 1C). In the P:E400:IF_occ_(LA) simulation both these distances consistently increased with time (Fig. 3 A,B). In the case of UP:E300:IF_occ_(LA) system, unlike in the P:E400:IF_occ_(LA), there was a drastic increase in both of these values during the first 100 ns and then gradually decreased with time, and slightly increased while the substrate was released into the cytoplasmic side (i.e., around 900 ns). However, final values of these two measures for this system were greater than the initial values (i.e., in the crystal structure). Overall, opening of the transporter was greater on the cytoplasmic side during the substrate release in P:E400:IF_occ_(LA) than in the UP:E300:IF_occ_(LA).

We also observed a salt bridge breaking on the cytoplasmic side (i.e., *D145 (TM5) - K417 (TM11)*) (Fig. 1C) in both the systems facilitating the transport (Fig. 3C).

Overall, our results indicate that the TM helices 5, 7, 10, and 11 play a key role in transportation of the substrate towards the cytoplasmic side after it binds in the substrate binding site. Helices 7, 10, and 11 belong to the C-domain and helix 5 belongs to the N-domain. Therefore, we conclude that C-domain dominates transportation process than the N-domain. This was also reflected in RMSF calculations, which indicate that the C-domain relatively fluctuates more than the N-domain in all the holo systems (Fig. S4 B,C). Particularly in the P:E400:IF_occ_(LA) system, region spanning residues 400-425 (loop between helices 10 and 11) fluctuates greater (≈8Å) (Fig. S4 C) compared to all other systems inferring that this region plays a key role in the transport process in this system. This also means that there are more interaction sites for the substrate in the C-domain than in the N-domain. On a side note, in the IF(Apo) systems N-domain fluctuated more than the C-domain, where as in the OCC(Apo) systems fluctuations of both domains were comparable (Fig.S4 A). OCC and IF forms majorly differ in the C-domain (spanning residues 400-425); this region fluctuates to a greater extent in the OCC than in the IF systems. The finding that increase in flexibility of the C-domain relative to the N-domain was not observed in the IF structures was rather interesting, as it may suggest that it was the interaction of substrate with C-domain that triggers the increased dynamics and not the other way around. This was in contrast to Fowler et al. report, which was based on short MD simulations (total 1.2*μs*) of IF(Apo) crystal structures, ^24^ that in both PepT_So_ and PepT_St_ dynamics of the C-domain dominates the N-domain. Such substrate-induced flexibility could perhaps be mechanistically relevant. *The asymmetric dynamics of the N- and C-domains observed in this study suggest that the PepT_St_ might not fit into the rocker and switch model, which was proposed earlier to describe functioning of the transporter*.^3,16^ It is possible that any single model may not completely explain the entire transport process in the POT transporters, particularly PepT_St_.

## Conclusions

In this study we investigated chemo-mechanical coupling involved in substrate/proton transport of the PepTSt transporter via microsecond level MD simulations. The key observations made in this study are that: (a) deprotonation/protonation of E300 is a must for the protein to bind and transport the substrate, (b) disengaging of residues E299 and E400 from interacting with the substrate promotes the release of substrate towards the cytoplasmic side, and (c) interaction of R33 with E299 and E300 acts as a molecular switch for the complete transition of protein from IF to OF state. Additionally, we identified that the C-domain was playing a major role in the functioning of the transporter, particularly TMs 7, 10, and 11, probably due to more interaction sites for the substrate on the C-domain compared to the N-domain. Finally, we report that the protein in all eight substrate bound systems (that differ in deprotonation/protonation of residues E299, E300 and E400) sampled different conformations from the crystal structure, indicating that there exists more physiologically relevant protonation states of the transporter apart from the ones studied here.

## Methods

12 different PepT_St_ systems (Table 1) were simulated via all-atom unbiased MD, each for 1 *μ*s. Three different crystal structures of PepT_St_ were utilized in this study: (a) PepT_St_ in IF state and complexed with HEPES (PDB: 6EIA), (2) PepT_St_ in IF_occ_ state and bound to Leu-Ala (LA) dipeptide (PDB: 5OXL), and (3) PepT_St_ in OCC state and bound with a phosphate ion in the substrate binding site (PDB: 5OXP)). ^22^ All three crystal structures were downloaded from the crystal data bank and processed using the MOE software. ^35^ Crystal waters and other components except dipeptide substrate were removed from the crystal structures, which resulted in one dipeptide bound PepT_St_ in IF_occ_ conformation, one PepT_St_ in IF(apo) conformation and one PepT_St_ in OCC(apo) conformation. The missing residues in these three structures were modeled using the MOE software and protonation states were assigned using the protonate 3D facility in MOE. These three PepT_St_ systems were modeled with either E300 protonated(P:E300:IF_occ_(LA)) or deprotonated(UP:E300:IF_occ_(LA)), resulting in six different systems. The substrate bound form was also modeled in six other protonation states: (1) E299 protonated (P:E299:IF_occ_(LA)), (2) E400 protonated (P:E400:IF_occ_(LA)), E299 and E300 protonated (P:E299,E300:IF_occ_(LA)), (4) E299 and E400 protonated (P:E299,E400:IF_occ_(LA)), (5) E299 and E400 protonated (P:E299,E400:IF_occ_(LA)), (6) E299, E300 and E400 protonated (P:E299,E300,E400:IF_occ_(LA)). OPM web server was used to orient protein in the membrane.^36^ Further, all 12 systems were built for MD simulations using the CHARMM-GUI web server. ^37^ Each system consists of one PepT_St_, ≈ 18,353 TIP3P waters,^38^ ≈ 280 POPE lipids, and 0.15M NaCl; and an additional dipeptide in the substrate bound systems. The size of the each system was ≈100×100×101 Å^3^, and contains ≈97,500 atoms. Each system was first energy minimized via conjugate gradient algorithm^39^ for 10,000 steps and then relaxed in multiple steps (see reference^40^ for details) for a total of ≈1.5 ns in NVT ensemble. Further, production simulations were started from the relaxed models, and data was collected every 0.5 ns for a total of 1 *μ*s in each case in an NPT ensemble. Simulations were conducted using NAMD 2.12^41^ in periodic boundary conditions and data analysis was conducted using various VMD plugins. ^42^ All components were modeled using CHARMM36 all-atom additive force field.^43^ A 2 fs timestep was used and simulations were carried at 310 K using Langevin integrator with a damping coefficient of *γ* =0.5 ps^−1^ and pressure was maintained at 1 atm using the Nosé-Hoover Langevin piston method.^44,45^ The non-bonded interaction cut-off distance was set to 10−12 Å, and particle mesh Ewald (PME) method^46^ was used to compute the long-range electrostatic interactions. We have used 24,000 data points (12 systems × 2 data points/ns × 1 *μs* = 24,000) for analysis in this work.

We define helices (TM) and loops (L) as follows: helix TM1: residues 13 to 46; TM2: 53 to 81; TM3: 83 to 104; TM4: 107 to 133; TM5: 142 to 173; TM6: 175 to 197; TMA: 213 to 242; TMB: 245 to 266; TM7: 282 to 312; TM8: 321 to 347; TM9: 354 to 383; TM10: 385 to^*a*^ Occluded PepT_St_ in apo form; ^*b*^ Inward-facing PepT_St_ in apo form. ^*c*^ Inward-facing Occluded PepT_St_ bound with substrate LA. UP and P refers to deprotonated and protonated residues.

## Acknowledgement

This research is supported by National Science Foundation grant CHE 1945465 and the Arkansas Biosciences Institute. This research is part of the Blue Waters sustained-petascale computing project, which is supported by the National Science Foundation (awards OCI-0725070 and ACI-1238993) and the state of Illinois. This work also used the Extreme Science and Engineering Discovery Environment (allocation MCB150129), which is supported by National Science Foundation grant number ACI-1548562. This research is also supported by the Arkansas High Performance Computing Center, which is funded through multiple National Science Foundation grants and the Arkansas Economic Development Commission.

## References

(1) Immadisetty, K.; Hettige, J.; Moradi, M. The Journal of Physical Chemistry B 2017, 121, 3644–3656.

(2) Ito, K.; Hikida, A.; Kawai, S.; Lan, V. T. T.; Motoyama, T.; Kitagawa, S.; Yoshikawa, Y.; Kato, R.; Kawarasaki, Y. Nature communications 2013, 4, 1–10.

(3) Lyons, J. A.; Parker, J. L.; Solcan, N.; Brinth, A.; Li, D.; Shah, S. T.; Caffrey, M.; Newstead, S. EMBO reports 2014, 15, 886–893.

(4) Newstead, S. Current opinion in structural biology 2017, 45, 17–24.

(5) Parker, J. L.; Mindell, J. A.; Newstead, S. Elife 2014, 3, e04273.

(6) Prabhala, B. K.; Aduri, N. G.; Hald, H.; Mirza, O. International journal of peptide research and therapeutics 2015, 21, 1–6.

(7) Guettou, F.; Quistgaard, E. M.; Tresaugues, L.; Moberg, P.; Jegerschöld, C.; Zhu, L.; Jong, A. J. O.; Nordlund, P.; Löw, C. EMBO reports 2013, 14, 804–810.

(8) Jensen, J. M.; Simonsen, F. C.; Mastali, A.; Hald, H.; Lillebro, I.; Diness, F.; Olsen, L.; Mirza, O. Peptides 2012, 38, 89–93.

(9) Newstead, S. Biochimica Et Biophysica Acta (BBA)-General Subjects 2015, 1850, 488–499.

(10) Smith, D. E.; Clémençon, B.; Hediger, M. A. Molecular aspects of medicine 2013, 34, 323–336.

(11) Meredith, D.; A, B. C. Cellular and Molecular Life Sciences CMLS 2000, 57, 754–778.

(12) Rautio, J.; Kumpulainen, H.; Heimbach, T.; Oliyai, R.; Oh, D.; Jarvinen, T.; Savolainen, J. Nature Reviews Drug Discovery 2008, 7, 255–270.

(13) Brandsch, M. Current Opinion in Pharmacology 2013, 13, 881–887.

(14) Anderson, C. M.; Thwaites, D. T. Physiology 2010, 25, 364–377.

(15) Jung, D.; Dorr, A. The Journal of Clinical Pharmacology 1999, 39, 800–804.

(16) Newstead, S.; Drew, D.; Cameron, A. D.; Postis, V. L.; Xia, X.; Fowler, P. W.; Ingram, J. C.; Carpenter, E. P.; Sansom, M. S.; McPherson, M. J.; Baldwin, S. A.; Iwata, S. 2011, 30, 417–426.

(17) Solcan, N.; Kwok, J.; Fowler, P. W.; Cameron, A. D.; Drew, D.; Iwata, S.; Newstead, S. The EMBO journal 2012, 31, 3411–3421.

(18) Molledo, M. M.; Quistgaard, E. M.; Flayhan, A.; Pieprzyk, J.; Low, C. Structure 2018, 26, 467–476.

(19) Doki, S.; Kato, H. E.; Solcan, N.; Iwaki, M.; Koyama, M.; Hattori, M.; Iwase, N.; Tsukazaki, T.; Sugita, Y.; Kandori, H.; Newstead, S.; Ishitani, R.; Nureki, O. Proceedings of the National Academy of Sciences 2013, 110, 11343–11348.

(20) Jardetzky, O. Nature 1966, 211, 969–970.

(21) others,, et al. Journal of the American Chemical Society 2019, 141, 2404–2412.

(22) Martinez Molledo, M.; Quistgaard, E. M.; Löw, C. FEBS letters 2018, 592, 3239–3247.

(23) Aduri, N. G.; Prabhala, B. K.; Ernst, H. A.; Jørgensen, F. S.; Olsen, L.; Mirza, O. Journal of Biological Chemistry 2015, 290, 29931–29940.

(24) Fowler, P. W.; Orwick-Rydmark, M.; Radestock, S.; Solcan, N.; Dijkman, P. M.; Lyons, J. A.; Kwok, J.; Caffrey, M.; Watts, A.; Forrest, L.; S, N. Structure 2015, 23, 290–301.

(25) Radestock, S.; Forrest, L. R. Journal of molecular biology 2011, 407, 698–715.

(26) Forrest, L. R. Science 2013, 339, 399–401.

(27) Quistgaard, E. M.; Molledo, M. M.; Löw, C. PloS one 2017, 12, e0173126.

(28) Parker, J. L.; Li, C.; Brinth, A.; Wang, Z.; Vogeley, L.; Solcan, N.; Ledderboge-Vucinic, G.; Swanson, J. M.; Caffrey, M.; Voth, G. A. Proceedings of the National Academy of Sciences 2017, 114, 13182–13187.

(29) Samsudin, F.; Parker, J. L.; Sansom, M. S.; Newstead, S.; Fowler, P. W. Cell chemical biology 2016, 23, 299–309.

(30) Immadisetty, K.; D Madura, J. Current Computer-Aided Drug Design 2013, 9, 556–568.

(31) Immadisetty, K.; Hettige, J.; Moradi, M. ACS Central Science 2019, 5, 43–56.

(32) Ogden, D.; Immadisetty, K.; Moradi, M. bioRxiv 2019, 708289.

(33) Harkey, T.; Kumar, V. G.; Hettige, J.; Tabari, S. H.; Immadisetty, K.; Moradi, M. Scientific reports 2019, 9, 1–12.

(34) Immadisetty, K.; Polasa, A.; Shelton, R.; Moradi, M. bioRxiv 2019, 707794.

(35) Chemical Computing Group Inc., Molecular Operating Environment(MOE), 2013.08 2016.

(36) Lomize, M. A.; Pogozheva, I. D.; Joo, H.; Mosberg, H. I.; Lomize, A. L. Nucleic acids research 2012, 40, D370–D376.

(37) Lee, J. et al. Journal of chemical theory and computation 2016, 12, 405–413.

(38) Jorgensen, W. L.; Chandrasekhar, J.; Madura, J. D.; Impey, R. W.; Klein, M. L. The Journal of chemical physics 1983, 79, 926–935.

(39) Reid, J. K. In Large Sparse Sets of Linear Equations; Reid, J. K., Ed.; Academic Press: London, 1971; pp 231–254.

(40) Jo, S.; Kim, T.; Im, W. PloS one 2007, 2, e880.

(41) Phillips, J. C.; Braun, R.; Wang, W.; Gumbart, J.; Tajkhorshid, E.; Villa, E.; Chipot, C.; Skeel, R. D.; Kale, L.; Schulten, K. Journal of computational chemistry 2005, 26, 1781–1802.

(42) Humphrey, W.; Dalke, A.; Schulten, K. Journal of molecular graphics 1996, 14, 33–38.

(43) Klauda, J. B.; Venable, R. M.; Freites, J. A.; O’Connor, J. W.; Tobias, D. J.; Mondragon-Ramirez, C.; Vorobyov, I.; MacKerell Jr, A. D.; Pastor, R. W. The journal of physical chemistry B 2010, 114, 7830–7843.

(44) Martyna, G. J.; Tobias, D. J.; Klein, M. L. The Journal of chemical physics 1994, 101, 4177–4189.

(45) Feller, S. E.; Zhang, Y.; Pastor, R. W.; Brooks, B. R. The Journal of chemical physics 1995, 103, 4613–4621.

(46) Darden, T.; York, D.; Pedersen, L. The Journal of chemical physics 1993, 98, 10089–10092.

