## Supporting Information for "Mechanistic picture for chemo-mechanical couplings in a bacterial proton-coupled oligopeptide transporter from Streptococcus thermophilus"

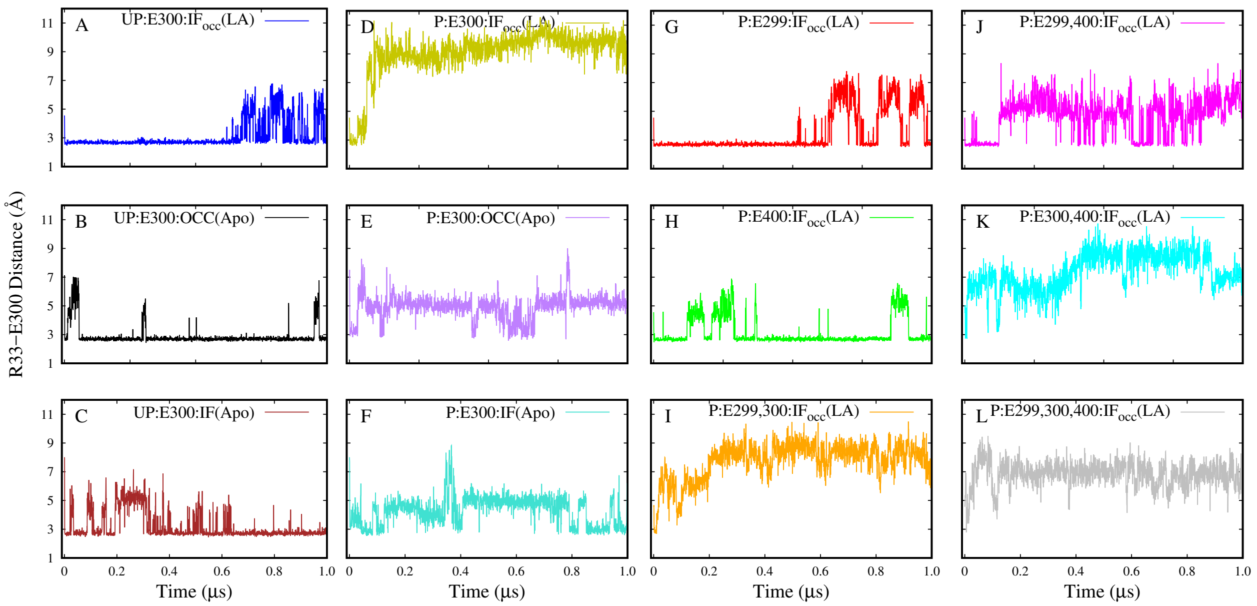


Figure S1: Time series of R33-E300 salt bridge distance. The presence of substrate destabilized this salt bridge in all the systems with E300 deprotonated(**A, G, H, J**), where as the absence of it kept this interaction intact (**B, C**). This salt bridge interaction was lost in all the E300 deprotonated systems as expected (**D, E, F, I, K, L**), although in the apo systems (**E, F**) the distance was smaller compared to the substrate bound systems.

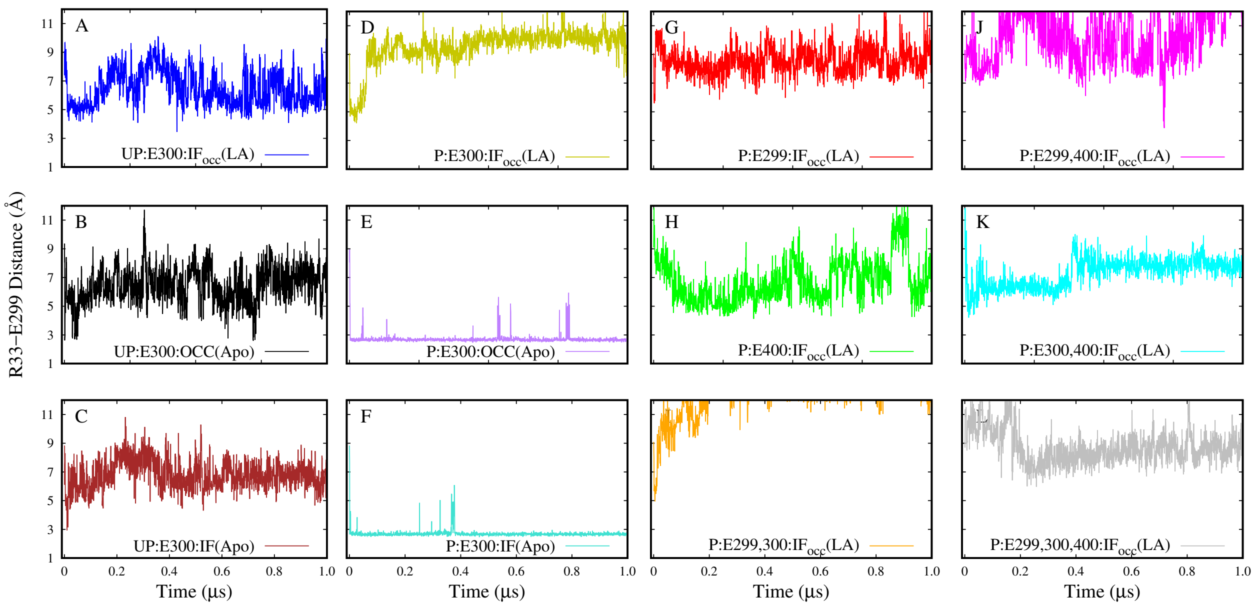


Figure S2: Time series of R33-E299 distance. Except in the P:E300:OCC(Apo)(**E**) and P:E300:IF(Apo)(**F**), this interaction was absent in all other systems.

Table S1: Substrate and protein contact frequency.

| **Systems** | | | | | | | | | | |
|---|---|---|---|---|---|---|---|---|---|---|
| E300:UP | **W296 87** | **Y68 76** | E299 69 | W427 62 | F428 53 | R26 51 | E400 45 | E300 38 | S431 37 | K126 32 |
| E300:P | **E299 98** | **E300 98** | **R33 97** | **S303 91** | **V304 81** | **M60 80** | **A64 78** | E400 66 | N328 59 | A61 55 |
| E299:P | **E400 98** | **V152 84** | **I332 81** | **R26 76** | **S404 75** | Y30 62 | N156 56 | N328 52 | Y29 46 | W296 39 |
| E400:P | **E299 90** | **W296 84** | E400 67 | W427 60 | I403 57 | F295 51 | S404 46 | F357 40 | A292 36 | I332 26 |
| E299E300:P | **R33 98** | **R26 92** | **Y68 71** | E400 65 | Y30 58 | N328 56 | W427 52 | P329 50 | I332 50 | V67 33 |
| E299E400:P | **W427 81** | **W296 77** | Y68 66 | F428 60 | S431 59 | R26 50 | E300 32 | E400 32 | I332 25 | Y30 25 |
| E300E400:P | **E299 98** | **R33 98** | **Y68 94** | W427 68 | W296 65 | R26 64 | E400 57 | K126 52 | V67 49 | T122 44 |
| E299E300E400:P | **R33 99** | **Y30 92** | **Y68 79** | P329 66 | E400 63 | A159 62 | I332 53 | W296 52 | W427 41 | I155 39 |

Contact frequencies of residues within 4 Åof LA were estimated. All probabilities were shown in percentage. Only top 10 interacting residues were shown in each case. LA interact with more residues in protonated than in the deprotonated systems. Particularly, this was difference was observed in the UP:E300:IF$_{occ}$(LA) and P:E300:IF$_{occ}$(LA) systems. Residues belonging to the helices 1, 7, 10 and 11 were predominantly interacting with the LA. Rarely residues of helices 2, 4 and 5 were also interacting with the LA. When E300 was protonated, LA prefers to interact with the residues of the N-domain (such as R26, Y30, R33, Y68).

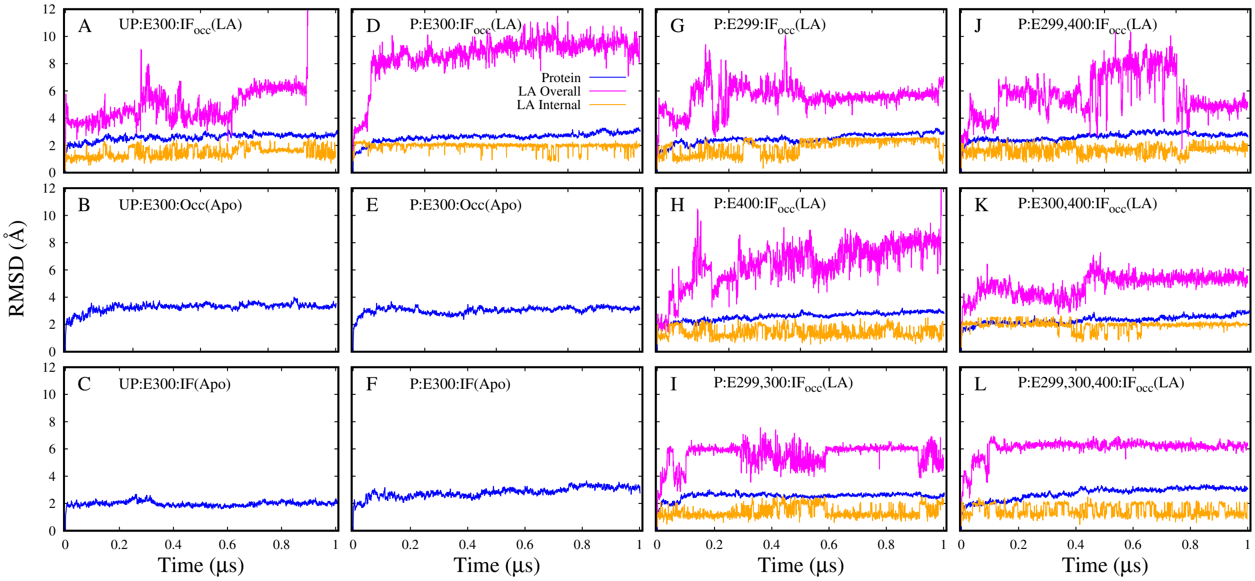


Figure S3: RMSD time series. Protein backbone, substrate overall, and substrate internal RMSDs were colored blue, magenta, and orange, respectively. Protein backbone RMSD of all systems stabilized around 3.5 Å. Internal RMSDs have not changed much in any case, indicating no conformational changes occurred in the substrate. Overall RMSD of substrates in E300 protonated systems (**D, I, K, L**) were relatively more stable than in the E300 deprotonated systems.

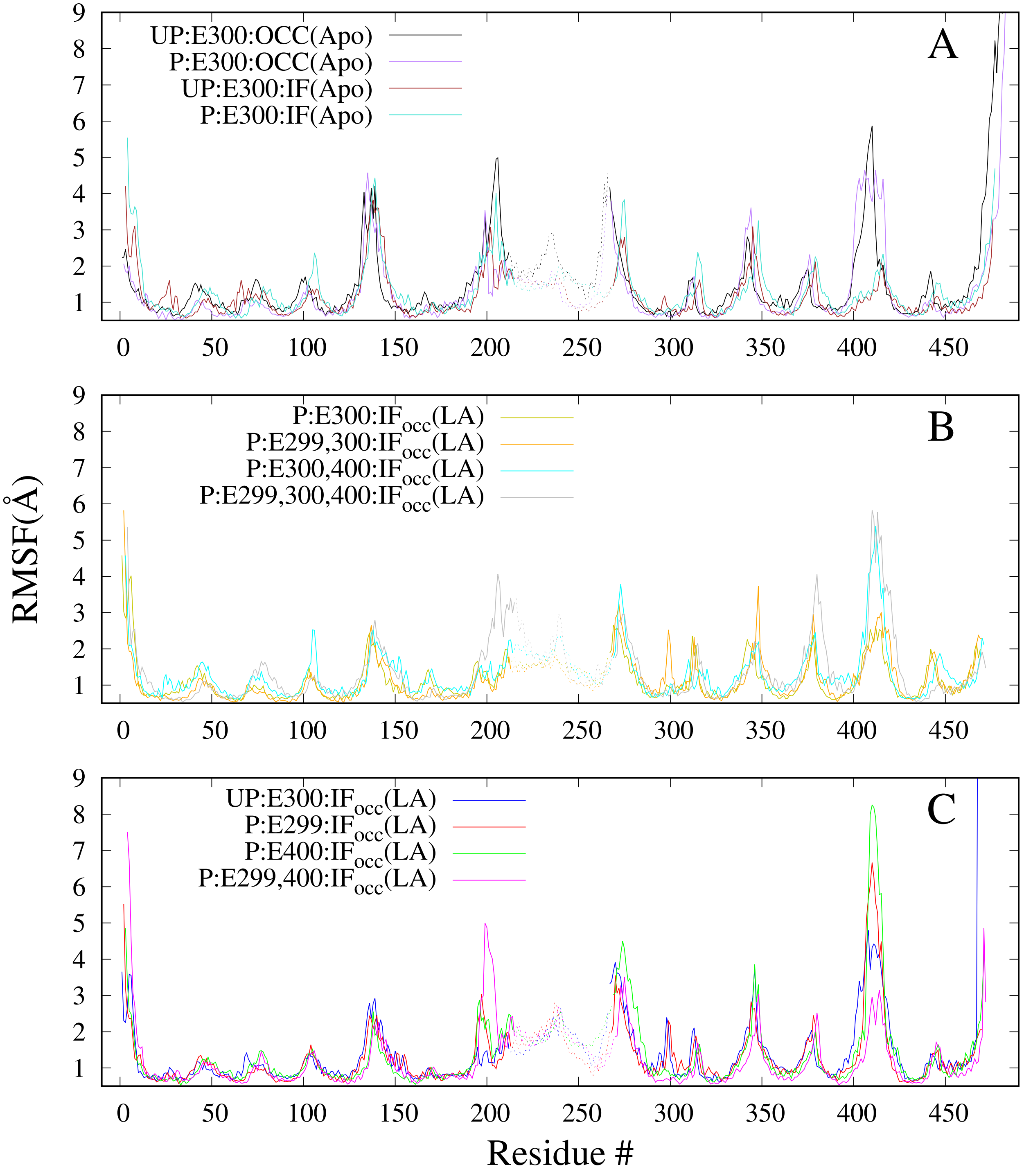


Figure S4: RMSF vs. Residue Number. All apo systems are shown in **A**, all E300 protonated systems in **B**, and the rest in **C**.

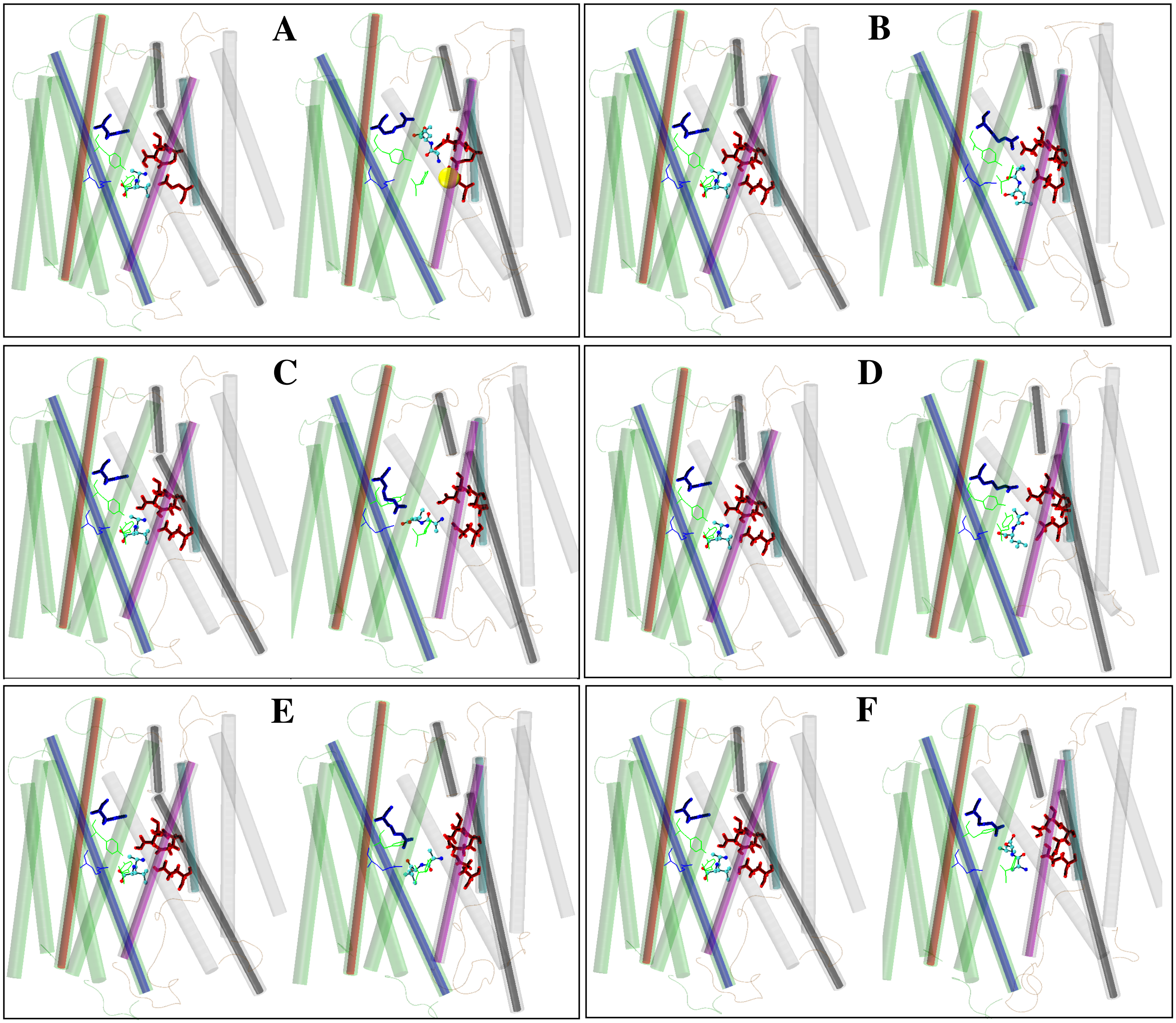


Figure S5: Substrate orientations in different protonation states. **A** is P:E300:IF$_{occ}$(LA), **B** is P:E299:IF$_{occ}$(LA), **C** is P:E299,E300:IF$_{occ}$(LA), **D** is P:E299,E400:IF$_{occ}$(LA), **E** is P:E300,E400:IF$_{occ}$(LA) and **F** is P:E300,E400:IF$_{occ}$(LA). Left and right snapshots in each panel represent starting and final orientations of the substrate.

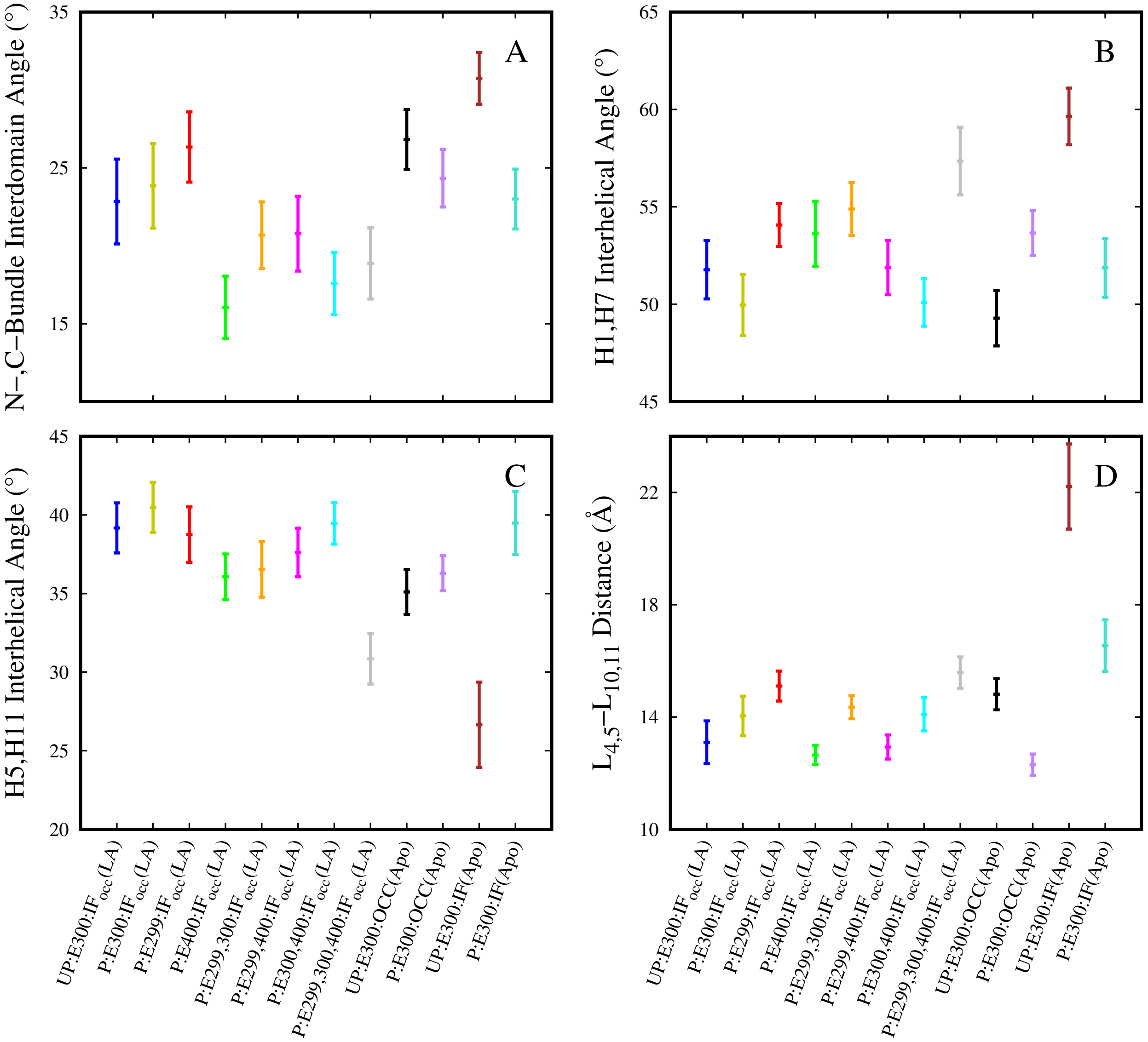


Figure S6: Impact of deprotonation/protonation on protein. Major impact of protonation of E300 was observed in the case of IF(Apo) systems. Last 100 of the simulation trajectory was considered for these calculations.
